# ARUNA: Slice-based self-supervised imputation for upscaling DNA methylation sequencing assays

**DOI:** 10.64898/2026.01.29.702677

**Authors:** Janmajay Singh, Wei-Hao Lee, Grace Yu, Vicky Yao

**Affiliations:** Department of Computer Science, Rice University 6100 Main Street, Houston, TX 77005, USA; Systems, Synthetic, and Physical Biology Graduate Program, Rice University 6100 Main Street, Houston, TX 77005, USA; Ken Kennedy Institute, Rice University 6100 Main Street, Houston, TX 77005, USA; Rice Synthetic Biology Institute, Rice University 6100 Main Street, Houston, TX 77005, USA

## Abstract

Whole-genome bisulfite sequencing (WGBS) can provide near-comprehensive, base-resolution maps of DNA methylation, transforming our understanding of epigenetic regulation in development and disease, but its cost is often prohibitive for many studies. Reduced representation bisulfite sequencing (RRBS) offers a cost-effective alternative that profiles a CpG-enriched subset of the genome at base resolution. Similar sequencing protocols for both assays pose an opportunity for cross-assay integration, presenting an opportunity for massively increasing sample sizes at whole-genome resolution. However, existing imputation methods are designed for within-assay scenarios and cannot handle the substantial CpG coverage differences between WGBS and RRBS. We introduce ARUNA, a self-supervised denoising convolutional autoencoder that predicts genome-wide CpG-level methylation using only a small subset of observed methylation values and CpG coordinates. By modeling methylation “slices,” spatially stacked windows that preserve local correlation structure, ARUNA captures biologically meaningful covariation while avoiding representation collapse. In simulation studies using the GTEx dataset, ARUNA successfully upscales RRBS-scale sparse methylomes (80-95% missingness) to whole-genome resolution, consistently outperforming baselines and maintaining robust performance across donor and tissue holdouts. When applied to real RRBS data from the ENCODE dataset, ARUNA outperformed state-of-the-art methods, with performance validated by matching upscaled RRBS samples to isogenic WGBS replicates. Source code for ARUNA can be found at https://github.com/ylaboratory/ARUNA.

## 1. Introduction

DNA methylation at cytosine-guanine (CpG) dinucleotides is a central epigenetic mechanism involved in cellular identity, gene regulation, embryonic development, and stem cell differentiation [1]. Dysregulation of DNA methylation has been implicated in many diseases, such as cancer, developmental disorders, neurological conditions, and autoimmune diseases [2, 3]. Although DNA methylation is one of the few epigenetic marks known to be mitotically heritable [4], it is also dynamic, changing with age, environmental exposures, and pathological states [5, 6]. Because these changes can be subtle and highly context-dependent, many discovery-driven studies aim to measure methylation at single-base resolution across the genome to capture its full regulatory complexity [7].

Several assays have been developed to measure DNA methylation, but bisulfite sequencing remains the only approach for whole-genome, base-resolution profiling. The two primary bisulfite sequencing methods differ substantially in cost and resolution: Whole-Genome Bisulfite Sequencing (WGBS) [8] provides comprehensive coverage (*>*75% of CpGs) but at high cost, while Reduced Representation Bisulfite Sequencing (RRBS) offers an economical alternative by using restriction enzymes to enrich for CpG-rich regions, covering only 10-20% of CpGs [9]. Apart from the initial target selection step in RRBS, both methods share nearly identical protocols involving bisulfite conversion and next-generation sequencing, and are therefore susceptible to similar noise profiles from bisulfite conversion errors and PCR bias. These similar technical characteristics make RRBS-to-WGBS upscaling and integration particularly attractive. Moreover, since RRBS introduces structured rather than random sparsity that favors CpG-dense regions [10], computational and machine learning methods can be used to leverage missingness patterns and bridge the gap in CpG coverage.

At the same time, large public repositories now contain thousands of high-quality WGBS samples across diverse tissues and conditions (e.g., GTEx [11], ENCODE [12], SRA [13]). These datasets encode rich information about local methylation structure and cross-sample covariance, suggesting the possibility of learning transferable genome-wide methylation patterns from public data and applying them to new assays. Successfully upscaling RRBS to whole-genome resolution would substantially increase effective sample sizes at genome-wide coverage, improving statistical power for downstream analyses [14] such as differentially methylated region detection, and faciliating analyses that benefit from comprehensive genomic coverage, such as methylation quantitative trait loci discovery. Importantly, while raw sequencing reads are often more difficult to access directly due to privacy concerns, processed data in standardized forms (e.g., BED files containing CpG coordinates and methylation beta values) are widely shared. In this work, we treat RRBS-to-WGBS upscaling as an imputation problem defined at the BED-file level and investigate whether models trained on public WGBS data can accurately recover genome-wide methylation maps from extremely sparse inputs, potentially enabling substantially more cost-effective whole-genome profiling.

Computational imputation of DNA methylation has been explored in several settings, but existing methods have important limitations for cross-assay upscaling. Classical machine learning approaches using random forests [15], Hidden Markov models [16], and XGBoost [17] rely heavily on engineered features and genomic annotations, requiring nontrivial preprocessing. More critically, these methods are designed to handle within-assay missingness and are not equipped to bridge the substantial genomic resolution gap between RRBS and WGBS. Imputation for single-cell bisulfite data has recently become an active area of research, employing Bayesian methods [18] and deep learning models [19], but these approaches focus on the sparsity patterns inherent in single cell technologies and not those seen in bulk WGBS or RRBS. Several “foundation models,” including CpGPT [20], MethylQUEEN [21], and MethylGPT [22], are pretrained on large collections of methylation data. However, most, if not all, of this data is drawn from array-based platforms that assay only a small subset of CpGs, limiting the models’ ability to generalize to true genome-wide imputation. Although these models aim for broad applicability, they typically do not support full-resolution predictions out of the box, and only CpGPT provides a fine-tuned model available for within-RRBS imputation. Crucially, no existing method provides genome-wide imputation from structurally sparse RRBS to WGBS resolution.

To address these limitations, we introduce ARUNA (**A**utoencoder for **R**econstruction and **U**pscaling of **N**oisy **A**ssays), a self-supervised framework designed for genome-wide methylation imputation across sequencing-based assays. ARUNA does not require raw sequencing reads, additional augmented data, or auxiliary feature engineering, instead operating directly on the standard BED format output that is the natural endpoint for most pipelines. The method is built around a novel slice-based representation, in which spatially aligned windows of CpGs are stacked across samples to capture both local methylation context and cross-sample covariance. Combined with a denoising convolutional autoencoder (DCAE) and CpG-wise positional encodings, this representation enables robust inference of genome-wide methylation patterns. We demonstrate ARUNA’s effectiveness using a combination of real cross-assay experiments and controlled generalization analyses. First, we establish real-world performance by training ARUNA on GTEx WGBS data and applying it to RRBS samples from ENCODE, verifying performance against isogenic WGBS replicates from the same ENCODE samples. We then perform complementary evaluations within the GTEx cohort to characterize robustness and generalizability, including donor holdout experiments to assess stability to inter-individual variability and tissue-holdout experiments to probe the model’s ability to capture biological variability across diverse tissue contexts. Across all scenarios, ARUNA consistently outperforms available methods, demonstrating accurate and generalizable genome-wide imputation.

## 2 Methods

### 2.1 Overview

Learning genome-wide DNA methylation structure from sequencing data presents several challenges that complicate imputation from sparse assays such as RRBS. First, methylation levels exhibit strong local correlation across nearby CpGs, leading to highly redundant, spatially structured features [23]. Second, inter-individual variability is relatively low within matched tissue/cell types, particularly in bulk tissue datasets [24, 25]. Third, available WGBS datasets typically contain far fewer samples than typically required for complex dense prediction tasks such as imputation, resulting in high-dimensional matrices with extreme feature correlation, a regime in which conventional machine learning methods often perform poorly or collapse to trivial solutions [26]. ARUNA addresses these challenges through three key design choices: (1) a novel slice-based representation which stacks spatially aligned genomic patches, augmenting training instances from limited data while allowing the model to jointly capture local co-methylation patterns and inter-individual covariance; (2) a self-supervised denoising objective trains the slice-based model to reconstruct artificially masked methylation values, preventing representation collapse and enabling the model to generalize to true missing data at inference time without paired RRBS-WGBS data; (3) a fully convolutional architecture exploits local CpG correlations through weight sharing and local receptive fields while maintaining a compact parameter space suitable for limited sample regimes.

ARUNA’s workflow comprises two main components: slice construction and self-supervised training (Fig. 1). Each sample methylome is first divided into contiguous genomic patches spanning 128 CpGs as the basic units for prediction and stacked across different samples to form slices. CpG-wise positional embeddings are concatenated with methylation values before being fed into the denoising convolutional autoencoder. During training, 90% of CpGs are masked at the slice level under a missing-completely-at-random (MCAR) scheme, forming the model’s reconstruction targets. During both training and inference, multiple slices are constructed for each target sample-patch pair by repeatedly sampling different cross-sample contexts. During inference, final imputed values for each anchor patch are obtained by averaging predictions across its multiple slice reconstructions.

**Figure 1.**
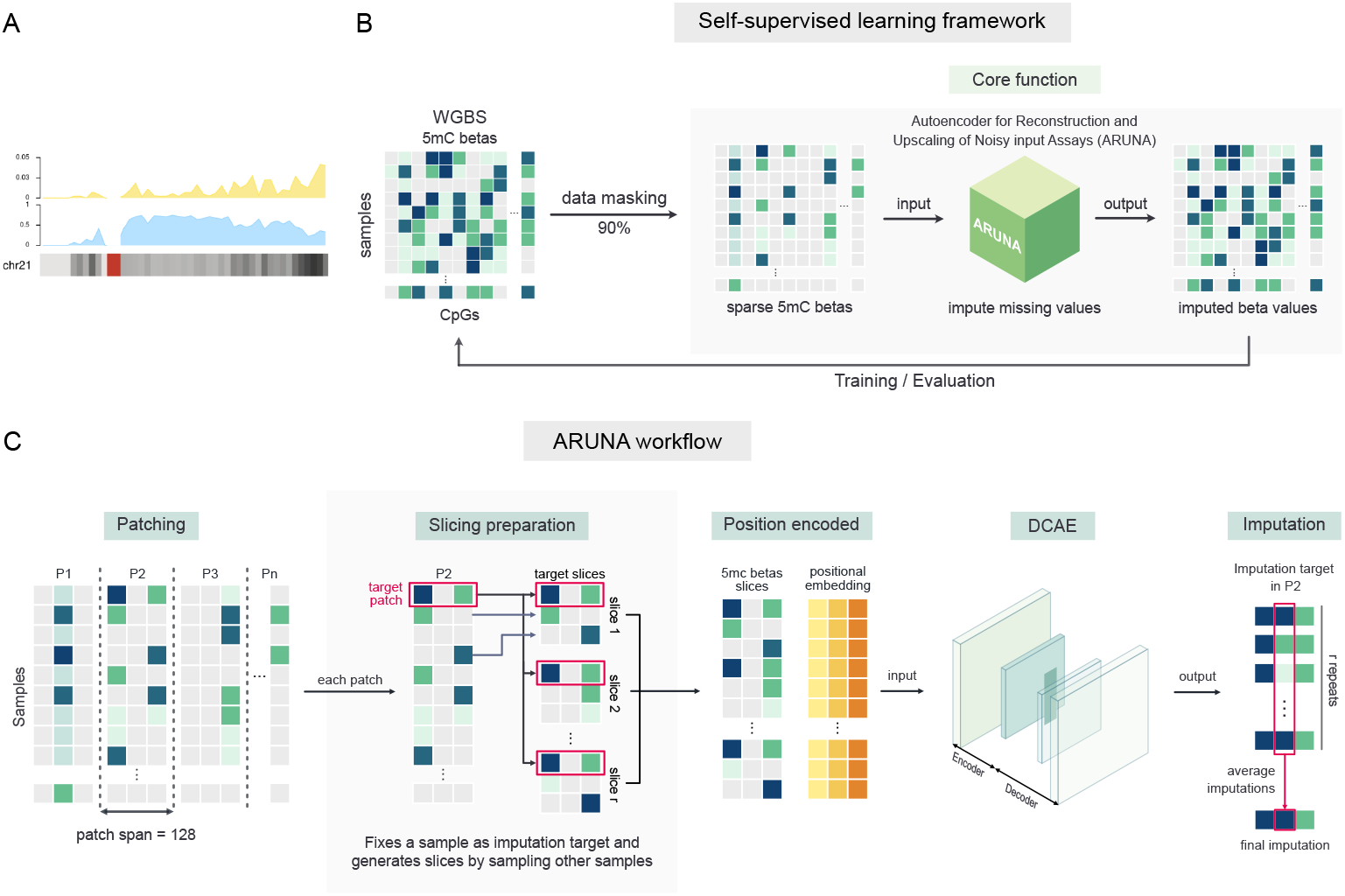
**(A)** Karyoplot of chr21 illustrating ARUNA’s motivating problem, showing CpG observation probabilities in (top) RRBS and (middle) WGBS, with reference CpG density overlaid along the chromosome (darker indicates higher density; centromere shown in red). **(B)** Overview of ARUNA’s self-supervised framework. Artificial masking is applied during training to generate a self-supervision signal, while during inference the sparse RRBS input is directly provided to the model for imputation. **(C)** ARUNA workflow detailing 5 steps: (i) Patch construction by partitioning each sample methylome into contiguous blocks of fixed size (128 CpGs); (ii) slice construction for a target sample–patch by randomly sampling a fixed number of supporting samples (slice height) and repeating this process to generate multiple “views” of the target; (iii) augmentation of beta values with positional embeddings encoding CpG genomic loci; (iv) input to the denoising convolutional autoencoder for slice-wise predictions; and (v) averaging *r* imputed outputs per target patch to produce the final prediction.

### 2.2 Slice-based representation

ARUNA’s central design choice is reformulating methylation data into “slices” as its fundamental training unit. By stacking patches at the same genomic position across multiple samples (slice “height” *h*), each slice provides both local co-methylation context and cross-sample variation. For each sample-patch, multiple slices are constructed by varying which samples are included (random sampling performed “repeat” times *r*). This design yields two critical advantages: (1) it enables efficient mini-batch gradient descent despite limited sample sizes, and (2) for a dataset of *n* samples, the number of potential slices scales combinatorially as 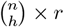, yielding a much larger pool of effective training units.

#### 2.2.1 Methylation Patches

Let a chromosome contain a fixed, canonical ordering of CpG sites indexed by ℳ = {1, 2, …, *M*}, where *M* is the total number of CpGs considered. Suppose we have *S* samples, and for each sample *s* ∈ {1, …, *S*} we denote its 1D methylome along this chromosome by 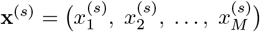, where 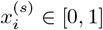 is the methylation level of CpG *i* in sample *s*.

Let *p* be a chosen patch size (number of CpGs per patch). We partition the CpG index set ℳ into contiguous, non-overlapping blocks of size *p*:

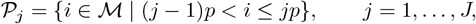

where *J* = ⌈*M/p*⌉ is the number of patches.

For sample *s*, the *j*-th patch is the length-*p* vector of methylation *β* values: 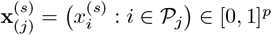 and thus each sample yields exactly *J* non-overlapping patches, and across all samples we obtain *S* × *J* patches. Padding is optionally applied to the last patch.

#### 2.2.2 Methylation Slices

To aggregate information across samples, we reorganize patch-level methylation vectors into fixed-height “slices.” Given a patch size *p*, each sample contributes a pool of *S* patch vectors 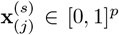 for patch index *j*. We specify a slice height *h*, which determines how many sample–patch vectors are stacked together. For each patch index *j*, a slice is an *h* × *p* matrix:

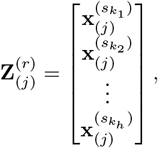

where the row indices 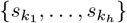 are drawn from the *S* − 1 samples (all but the target patch’s sample) with replacement, ensuring that slices consistently represent the same genomic interval across individuals. Slices do not mix different patch indices by construction. To increase diversity, we generate multiple independent slices per patch by repeating the sampling procedure *r* times, yielding alternative “views” of the same genomic window. In our experiments, we found an optimal slice dimension to be (8, 128), stacking 8 patches of 128 CpGs.

### 2.3 Learning Framework

#### 2.3.1 Self-supervision

To train the model to handle sparse inputs without requiring paired RRBS-WGBS data, we employ a denoising self-supervised approach. The model is trained on WGBS data in which additional missingness is artificially introduced according to our MCAR simulation framework, forcing it to learn robust methylation patterns that generalize to extreme sparse data (as found in RRBS) at inference time.

For each CpG *i* in a slice, let *x*_*i*_ denote the ground-truth methylation value and 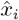 the model prediction. We associate two binary masks with each CpG: *c*_*i*_ ∈ {0, 1}, indicating whether the CpG was missing in the ground truth BED file (*c*_*i*_ = 1 if missing), and *m*_*i*_ ∈ {0, 1}, indicating whether the CpG was artificially masked during training (*m*_*i*_ = 1 if masked, typically with probability 0.9 in our experiments).

The positions that remain visible to the model in the input are tracked with a reconstruction mask *r*_*i*_ = (1 − *c*_*i*_)(1 − *m*_*i*_). Critically, these visible positions are not used for loss computation or evaluation. Instead, we train and evaluate only on CpGs that were originally observed but deliberately masked, using the training mask *t*_*i*_ = (1 − *c*_*i*_)*m*_*i*_.

This ensures the model learns to predict missing values from observed context rather than simply memorizing inputs. The self-supervised training objective is the masked mean squared error over all CpGs in the batch: 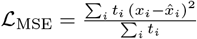.

#### 2.3.2 Denoising Convolutional Autoencoder

To effectively model spatial methylation relationships while learning across-individual beta-value variations, we chose a fully convolutional architecture for our model. In addition, we assumed that the true underlying biological variability would be low-dimensional relative to the number of CpGs, and decided to compress each input slice with a denoising autoencoder. The goal of the network is therefore to reconstruct the original slice *x* from its masked version 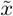.

The DCAE implements this mapping through a fully convolutional, symmetric pyramidal encoder-decoder architecture built using 2D CNN layers. The encoder *f*_*θ*_ applies a sequence of convolutional layers to extract a latent representation *z*:

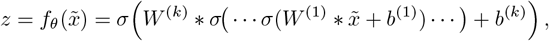

where *W* ^(*i*)^ and *b*^(*i*)^ are convolution kernels and biases, * denotes convolution, and *σ*(·) is a nonlinear activation (ReLU). The latent feature map *z* ∈ ℝ^*d*^ captures the local methylation structure learned from the noisy input. The decoder *g*_*ϕ*_ then mirrors the encoder with a sequence of transposed convolutions, reconstructing a full-length prediction 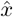. The overall DCAE therefore defines the mapping 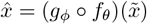, and was trained end-to-end to minimize the masked loss using pytorch [27]. More details on model architecture can be found in Supplementary Table 1.

#### 2.3.3 Positional encoding

CpG are unevenly distributed along the genome, such that a fixed-width patch may span loci separated by substantial genomic distances [28]. To expose this structure to the model, we augment each slice with an additional channel containing positional encodings for the CpGs in the patch. This channel encodes both the ordering of CpGs within a patch and their absolute genomic locations, providing biologically meaningful spatial context. We use a sinusoidal positional encoding scheme [29], adapted to DNA methylation data, to represent genomic position at multiple spatial scales. For a patch containing *p* CpGs, we construct a *p*-dimensional positional encoding vector for each CpG at genomic coordinate pos:

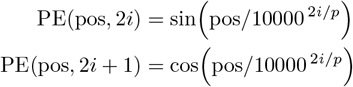

This construction assigns each CpG a continuous, multi-scale representation of its absolute genomic position. When concatenated with the methylation slice, the resulting input enables the model to distinguish CpGs that are nearby in index but distant in genomic space, while preserving relative structure within each patch.

### 2.4 Data Preprocessing and Missingness Simulations

Three datasets were used in this study: GTEx, ENCODE and, SRA. All 126 GTEx samples were WGBS and formed the primary training and evaluation cohort for our simulation studies. The samples spanned multiple tissue groups, including brain (*n* = 32), thyroid (*n* = 30), lung (*n* = 27), and a heterogeneous collection of additional tissues (*n* = 37). Both WGBS (*n* = 16) and RRBS (*n* = 17) data from ENCODE were used to assess across-dataset generalization and included both primary tissues and cell lines. In this study, we focused on ENCODE primary tissues: mammary epithelial cells and skeletal muscle myoblasts, excluding cancer cell lines. Each ENCODE sample was accompanied by an isogenic replicate, providing a high-confidence reference. Finally, 498 RRBS accessions from SRA were used to estimate empirical CpG-level coverage distributions for simulating RRBS-like missingness patterns.

#### 2.4.1 Uniform bioinformatics pipeline

All datasets were processed with standardized best-practice pipelines to avoid preprocessing-associated batch effects. WGBS samples were trimmed using `TrimGalore` [30], then `Bismark` [31] was used for alignment to the GRCh38 reference, deduplication, and methylation calling was performed followed by strand merging. RRBS samples were processed using analogous pipelines, with `TrimGalore` run in `–rrbs` mode and deduplication omitted in accordance with current best practices for RRBS libraries. For both assays, methylation calls were converted to BED-level CpG methylation matrices using a common canonical CpG set derived from the reference genome.

#### 2.4.2 BED-level missingness

We simulated missingness directly at the BED-file level and independently for each sample-CpG entry.

For MCAR experiments, we specified a target missingness level *ρ* as proportion of missing CpGs. Missingness was introduced by drawing *u*_*i*_ ∼ Uniform(0, 1) for each 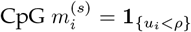 so that each CpG is masked with a fixed probability *ρ*.

To generate RRBS-like coverage patterns, we aggregated 498 RRBS accessions from SRA and computed CpG-wise empirical probabilities of observation *ρ*_*i*_ = Pr(*Obs*(*CpG*_*i*_)). These values were merged onto our canonical CpG set; CpGs absent from the SRA accessions were assigned *ρ*_*i*_ = 0. Missingness was then simulated using 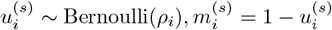, where 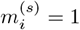 indicates a missing entry. This yielded nonuniform RRBS-like sparsity patterns, enriched in CpG islands.

### 2.5 Competing Methods

We compared ARUNA with several models, ranging from simple baselines to state-of-the-art imputation and foundation-model-based methods. The simple *Patch-Means* baseline imputes each CpG using the mean value within its patch. The patch-wise means were computed directly on the validation set. BoostMe [17] incorporates a sampleAvg feature by default, which would directly leak information from held-out samples in our evaluation regime. Usually, BoostMe trains one model per sample, and it counters leakage by enforcing a CpG-level or a chromosome-level holdout during training and inference. However, since we train one model per chromosome, the only way to ensure no leakage was to disable BoostMe’s sampleAvg feature. Still, the model trained on our test folds (unseen by ARUNA), giving BoostMe a significant advantage. Finally, we also compare against CpGPT, a large foundation model pretrained on a diverse and extensive collection of methylation datasets. The authors have released several task-specific fine-tuned variants, including a model optimized for intra-RRBS imputation. We used this pretrained and fine-tuned model as provided to generate imputed methylation values.

## 3. Results

Due to the novelty of our problem setting, none of our baselines were originally designed to impute from such extreme sparsity levels. ARUNA and simple statistical generate predictions at all CpG sites, whereas BoostMe fails to impute CpGs for which it cannot construct features, resulting in missing predictions for approximately 20% of CpGs. CpGPT, which was optimized for within-RRBS imputation, produces predictions only for CpGs present in its training vocabulary, resulting in a substantially more limited genomic coverage (∼2% of CpGs). Consequently, some evaluations are necessarily restricted to CpGs for which all methods provide predictions (limited primarily by CpGPT), though we focus on assessing near genome-wide performance based on the (∼80%) of CpGs BoostMe could predict, excluding CpGPT. Notably, when predicting on the stringent full intersection setting, this approximates a within-RRBS evaluation setup, since CpGPT selects for positions observed in RRBS data. Results were verified across chromosomes 1, 11, and 21 with all results here based on chr 21 for conciseness.

### 3.1 ARUNA enables genome-wide RRBS-WGBS upscaling on ENCODE data

We first evaluated ARUNA in a realistic cross-assay setting using matched RRBS and WGBS data from the ENCODE project. ENCODE provides isogenic replicates profiled using both assays, enabling direct validation of RRBS-to-WGBS upscaling against whole-genome ground truth. In this experiment, ARUNA was trained exclusively on GTEx WGBS samples and applied without retraining to ENCODE RRBS samples, reflecting a practical deployment scenario in which models trained on public reference cohorts are transferred to new datasets.

Across ENCODE samples, ARUNA achieved the lowest imputation error among all methods when evaluated on CpGs shared across models (Fig. 2A). Performance gains were observed across a broad range of CpG context, including CpG islands, shores, shelves, and open-sea regions, indicating that ARUNA does not rely on CpG density for accurate reconstruction. In contrast, CpGPT clearly performed better in more CpG-dense contexts (islands, shelves, shores) and significantly worse in open sea regions, consistent with its focus on RRBS-enriched regions [10]. BoostMe generally did not perform well on the restricted set of CpGs, oftentimes performing worse than the patch-means baseline.

**Figure 2.**
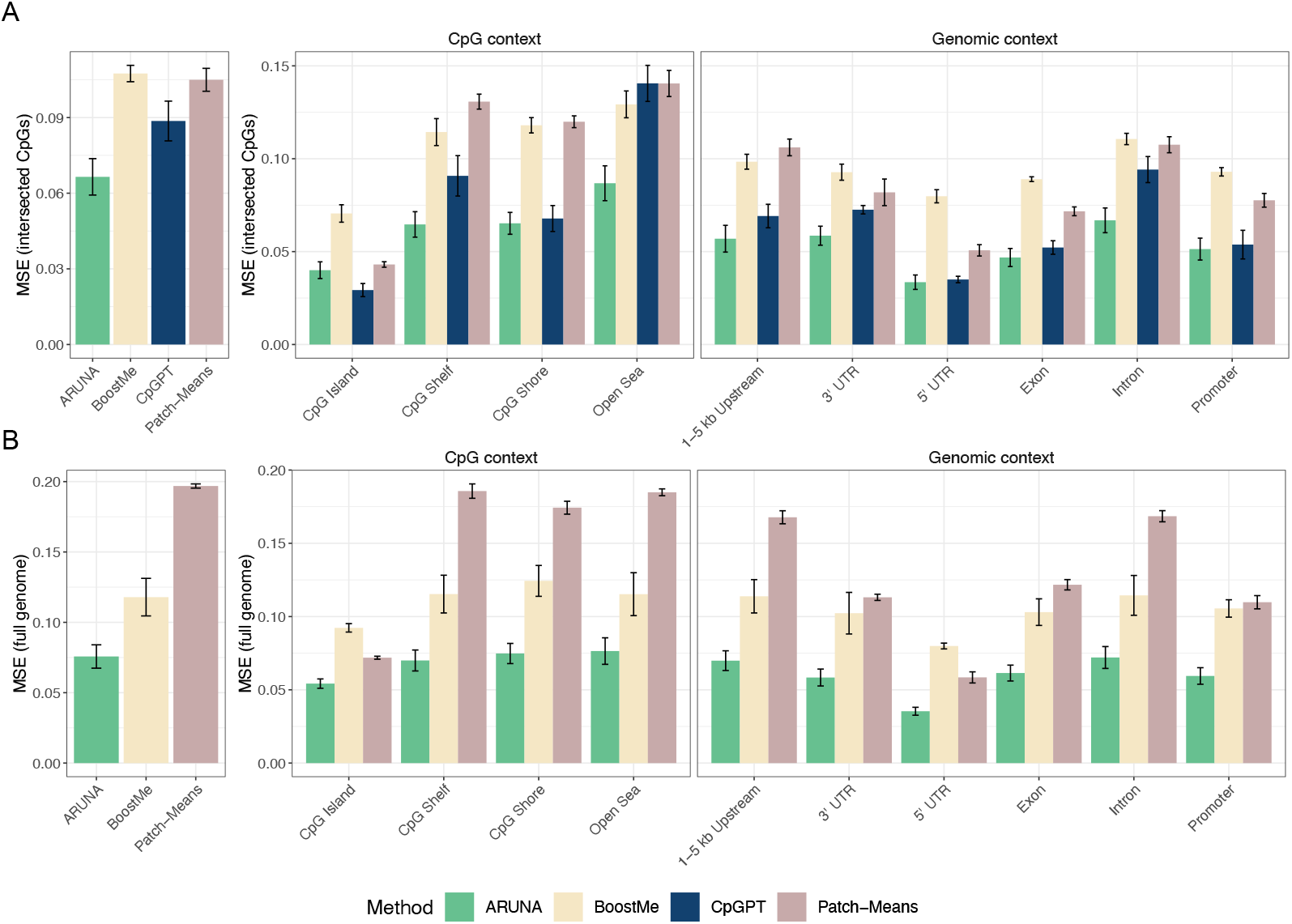
Comparative performance for the across-dataset generalization evaluations, with ARUNA trained on GTEx WGBS with simulated missingness and all models tested on ENCODE-RRBS. Matched ENCODE-WGBS samples were used as ground truth. Figures show MSE over test set samples genome wide (left), in CpG contexts (middle) and genic contexts (right). **(A)** FullIntersect (3, 888 *±* 401 CpGs); comprising mostly RRBS CpGs and **(B)** GenomeScale (400, 753 *±* 4972 CpGs) excludes CpGPT for broader evaluation across the methylome.

When evaluated at genome scale, BoostMe’s performance improved over patch-means, but ARUNA consistently had the strongest performance (Fig. 2B). This pattern was consistent overall, especially for CpG contexts, where ironically BoostMe performed particularly poorly on CpG islands, which would intuitively be the easiest to predict. In general, we find that ARUNA is capable of accurate genome-wide RRBS-to-WGBS upscaling in a real cross-assay setting, regardless of what collection of CpGs we analyze.

### 3.2 ARUNA robustly imputes genome-wide methylomes across held-out donors

We next evaluated ARUNA under a donor-holdout setting using GTEx data to assess inter-individual generalization. Donor identifiers from GTEx metadata were used to construct a 3-fold split in which all samples from a given donor were assigned to a single fold, ensuring that test donors were never observed during training. Donor representation was imbalanced (e.g., donor PT-12WSD contributed 15 samples, whereas 17 donors contributed a single sample each), while gender distribution was approximately balanced (66 female, 60 male samples). Importantly, this split did not induce tissue-level holdout: all tissues present in the validation set were also observed during training. Model training and evaluation were repeated across five random seeds. ARUNA was trained on MCAR-simulated data with a missing proportion of 0.9, and evaluation was performed on simulated RRBS data on the held-out fold.

To enable direct comparison across all models, we first evaluated performance on the intersection of CpGs for which predictions were available from every method. Within this shared CpG set, ARUNA achieved the lowest MSE, outperforming all other methods (Fig. 3, top). Similar to our findings for generalization between RRBS and WGBS data for ENCODE, CpGPT performed better in CpG dense regions. Even in CGIs, however, ARUNA achieved lower error (0.016 ± 0.003) than CpGPT (0.020 ± 0.003). For other methods, performance generally degraded from CGI to shelf, shore, and open-sea regions, indicating a dependence on CpG density. In contrast, ARUNA remained relatively invariant to this trend, likely due to its slice-based representation augmented with positional information, which enforces a constant number of CpGs per input unit. For non-CpG–centric genomic annotations, all models achieved their lowest errors in 5^*′*^ UTRs, consistent with known links between CpG density and methylation variability in the region [32] (Supplementary Figure 1).

**Figure 3.**
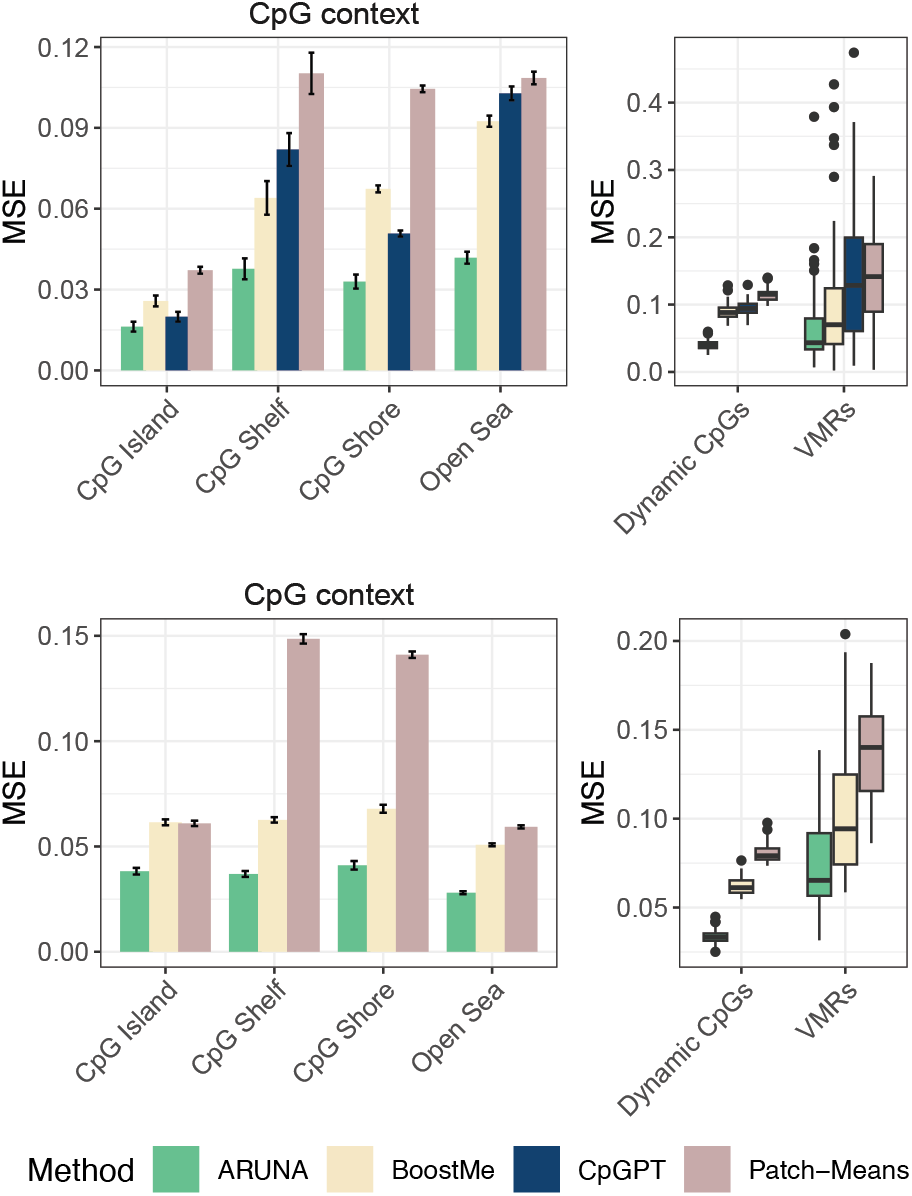
Performance evaluations for the donor-holdout simulation setup. Training and evaluation was performed using GTEx data with simulated missingness (mcar at 90% for training ARUNA and rrbs-like simulation for evaluations). Figures show MSE over held-out donor samples across aggregated over multiple folds and seeds stratified by CpG-context (left) and two additional contexts to better evaluate model performance on CpGs with high across-tissue (“Dynamic”) and across-donor (“VMRs”) variance (right). (Top) FullIntersect (2, 985 *±* 423 CpGs); (Bottom) GenomeScale (366, 887 *±* 4575 CpGs).

In addition, we evaluated performance on two subsets expected to be more challenging for imputation: Dynamic CpGs, which exhibit higher variability across tissues and individuals [24], and Variably Methylated Regions (VMRs), which are known to show high inter-individual variation [11]. For VMRs, we used the ubiquitous (non–tissue-specific) set defined in the original study. Restricting evaluation to the more challenging Dynamic CpG and VMR subsets further reduced the comparable CpG set to 973 ± 144 Dynamic CpGs and 5 ± 4 ubiquitous VMR CpGs. As expected, performance deteriorated for all models in these high-variability regions. Despite increased variance in VMR performance under donor holdout, ARUNA consistently outperformed baseline methods.

We further assessed calibration to evaluate imputation quality across the full range of methylation beta values (Fig. 4). ARUNA exhibited superior calibration, closely tracking the diagonal and faithfully capturing the bimodal distribution characteristic of DNA methylation, while maintaining accuracy at intermediate beta values. This behavior is particularly important, as intermediate methylation has been implicated as a conserved regulatory signature and associated with transcription regulation [33, 34]. ARUNA produced slightly conservative predictions in highly methylated regions, rarely predicting values exactly at 1, likely due to the use of a final sigmoid activation. BoostMe was well calibrated for hypomethylated CpGs but showed a bias toward hypermethylated predictions, while CpGPT largely failed to model intermediate methylation, instead collapsing predictions toward 0 or 1. This behavior may reflect CpGPT’s training on high-depth RRBS data and its bias toward CpG island–rich regions.

**Figure 4.**
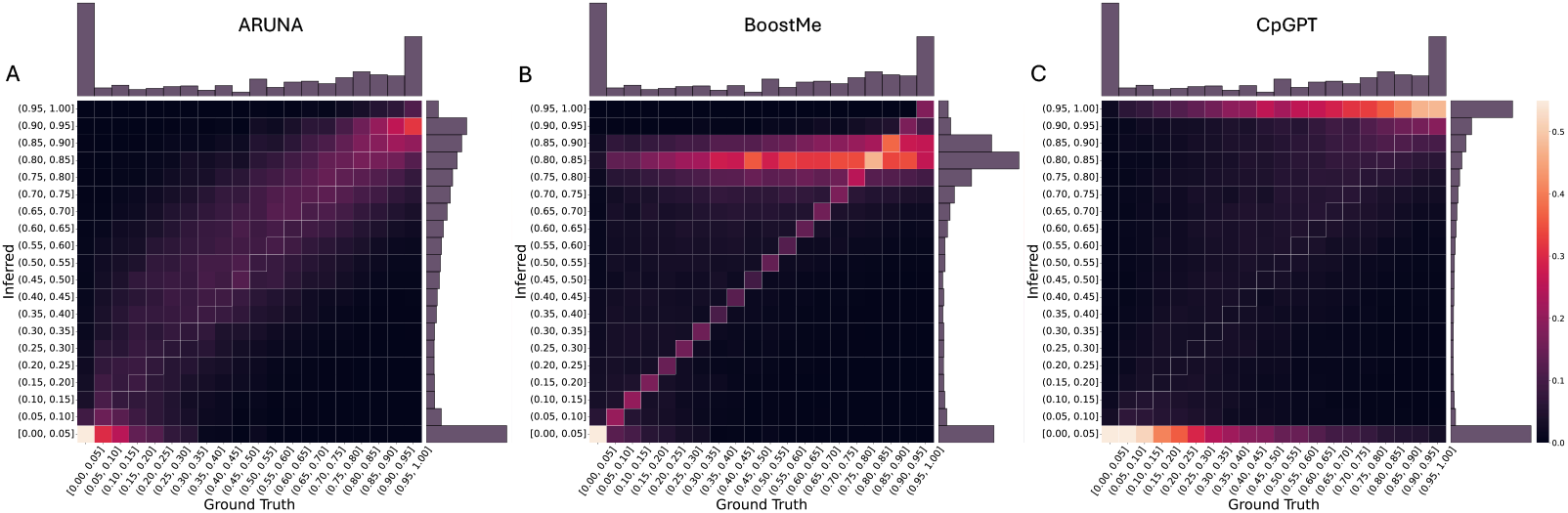
Model calibration across the full range of methylation beta values. Heatmaps show the relationship between true and inferred CpG beta values, with column-wise normalization indicating the relative proportion of CpGs compared to the true frequency within a beta value range. Marginal histograms depict the distributions of true and inferred CpG beta values, revealing the characteristic bimodal methylation pattern. The diagonal denotes perfect calibration and serves as a gold standard.

When evaluated genome-wide on all CpGs for which full-resolution predictions (Fig. 3, bottom), as expected, overall error increased for all models relative to the shared CpG evaluation. Nevertheless, ARUNA consistently achieved better performance across CpG contexts. Notably, ARUNA achieved substantially lower error on the Dynamic CpG set (95,367 ± 851 CpGs; MSE 0.033 vs. 0.062 for BoostMe) and showed only a minor degradation in performance on VMRs (158 ± 16 CpGs), indicating robustness to regions of elevated inter-individual variability.

To further probe performance in CpG-sparse settings, we analyzed imputation error as a function of the distance between a target CpG and its nearest observed neighbor (Fig. 5). As expected, performance degrades with increasing distance for both models. However, ARUNA consistently stabilizes at a lower error floor than BoostMe, maintaining strong performance even for CpGs lacking observations within a 100 kbp window. This suggests that ARUNA’s convolutional, slice-based representation captures broader methylome structure and enables modeling of long-range, non-local dependencies.

**Figure 5.**
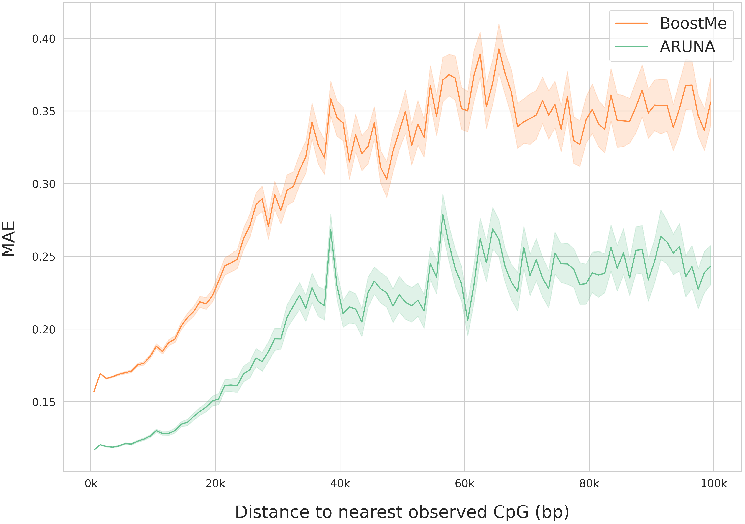
Imputation mean absolute error (MAE) seen against distance to the nearest observed CpG. Shaded regions indicate the 95% confidence interval for the estimated mean error. The plot is truncated at 100 kbp, beyond which model performance stabilizes.

In general, we see that ARUNA consistently outperforms existing imputation methods in a stringent donor-holdout setting, supporting its ability to generalize across individuals for genome-wide methylation upscaling and within-RRBS imputation.

### 3.3 ARUNA generalizes methylation upscaling to unseen tissues

We next examined ARUNA’s ability to generalize across tissues by performing tissue-holdout experiments in a leave-one-group-out setting. Using the GTEx dataset, tissues were grouped into four categories. All brain regions were aggregated into a single *Brain* holdout fold, with analogous groupings for *Thyroid* and *Lung*. The remaining tissues, which span diverse anatomical contexts but are each represented by relatively few samples, were grouped into a single *Other* fold, comprising skin, skeletal muscle, heart (left ventricle), pituitary, mammary tissue, and esophagus-muscularis. This grouping strategy was chosen to approximately balance the number of samples across the training and test folds, thereby reducing confounding effects arising from extreme disparities in train/test set sizes. Training and testing of the models were similar to the donor-holdout setup. We further included a cell-type-specific CpG subset from [35]. These are uniquely unmethylated loci identified in the original study from FACS-purified samples. Because these CpGs reflect strong cell-type specificity, they are well-suited for evaluating generalization to unseen tissues.

Again, we first evaluated performance on the subset of CpGs for which all methods produced predictions (Fig. 6, top). Across tissue holdout folds, ARUNA consistently achieved lower imputation error than other methods. Overall trends and relative performances remained similar to the donor-holdout setup, with slightly higher error rates across all models and regions, suggesting imputing for unseen tissues is a more difficult task. This is in line with known patterns since biological variation across tissues or cell types is stronger than inter-individual variation. Significant difference across folds was not observed, suggesting that with enough training samples, generalization to unseen tissues plateaus at a similar performance. Performance on the cell-type-specific loci was interesting to see since the average performance of both CpGPT and ARUNA was better on this subset than any other genomic region for all tissue holdout folds. Since in this setting, evaluation is limited to RRBS CpGs, this suggests CpGPT accurately imputes tissuespecific hypomethylation by leveraging long-distance relationships across its RRBS-specific CpG vocabulary. Interestingly, however, ARUNA still outperforms CpGPT on all but the skeletal muscle tissue (in the *Other* fold), suggesting ARUNA’s convolution architecture is capable of similar long-range modeling without the added model complexity. BoostMe’s average imputation performance remains close to that on other genomic regions, suggesting it can not capture these highly specific loci by simply modeling nearby CpGs methylation states.

**Figure 6.**
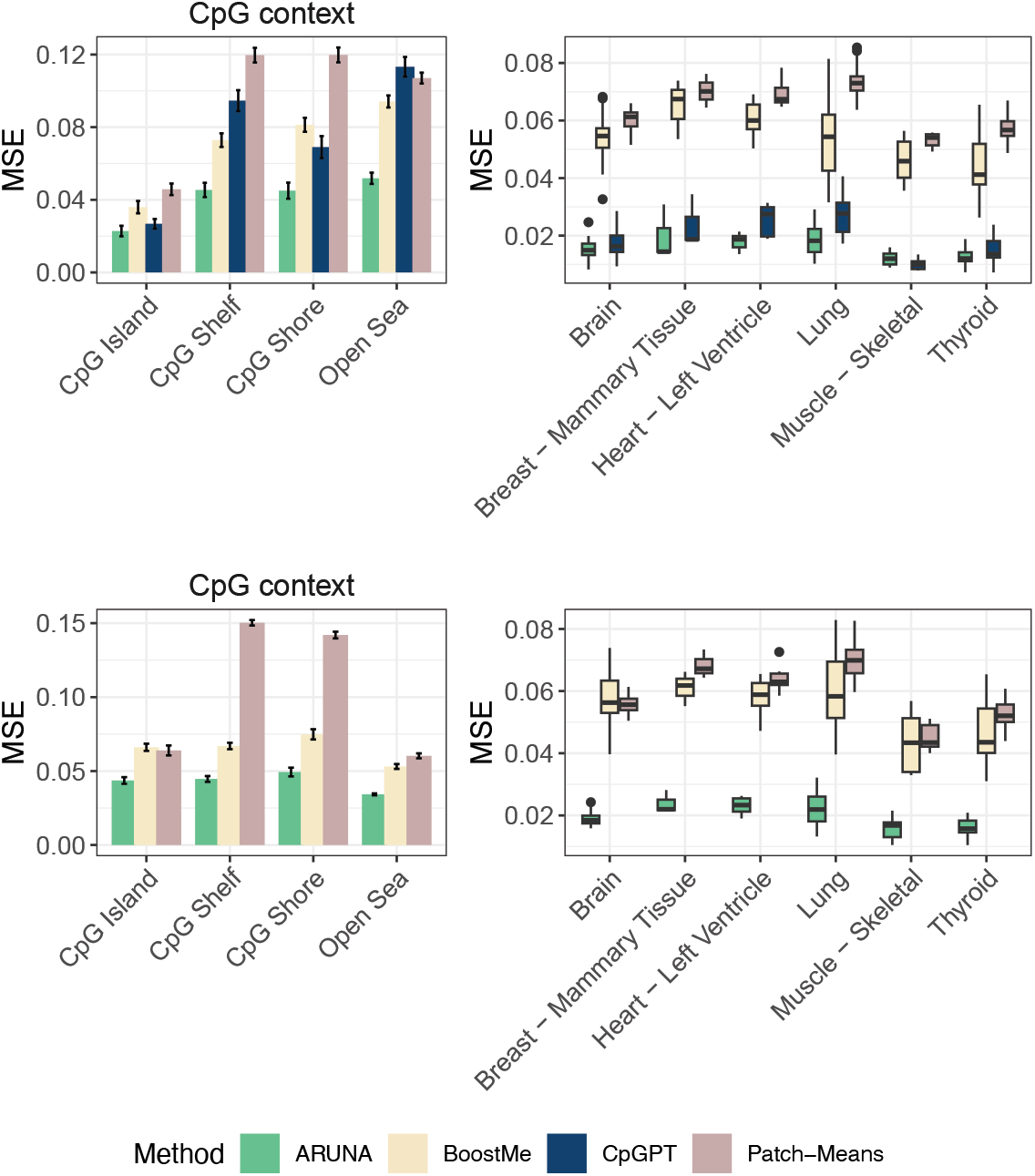
Performance evaluations for the tissue-holdout simulation setup. Generalization of models to unseen tissue types was assessed with MSE computed over the held-out tissue folds and seeds and stratified by CpG-context (left). Performance was also tested on a set of highly cell-type specific, unmethylated CpGs for each tissue group (right). (Top) FullIntersect (4,242 ± 1051 CpGs); (Bottom) GenomeScale (364,638 ± 5108 CpGs)

In addition to the shared CpG evaluation, we assessed performance at genome scale (Fig. 6, bottom), which again demonstrates that ARUNA is capable of maintaining stable, consistent performance across the genome, including across tissues. For highly cell type specific CpGs, BoostMe’s performance approximated that of the simple patch means baseline. We find that ARUNA captures methylation structure that generalizes across tissues, enabling robust imputation for tissue types not observed during training. Performance stratified by several genic contexts can be seen in Supplementary Figure 2.

## 4 Discussion

In this work, we introduced ARUNA, a slice-based self-supervised framework for genome-wide DNA methylation imputation and RRBS-to-WGBS upscaling directly from standard BED-format methylation matrices. ARUNA combines three key components: a slice representation that stacks spatially aligned CpG windows across samples to expose both local co-methylation structure and cross-sample variation; a denoising self-supervised objective that trains on WGBS without paired RRBS–WGBS data by training on artificially masked CpGs; and a fully convolutional autoencoder augmented with CpG-wise positional encodings to exploit local structure while remaining parameter-efficient in limited-sample regimes. Across real cross-assay evaluations on ENCODE RRBS with matched isogenic WGBS replicates and controlled simulations in GTEx WGBS, ARUNA consistently achieved the lowest error among available baselines and demonstrated robust generalization under donor and tissue holdouts. These results support ARUNA as a practical approach for converting structurally sparse, sequencing-based methylomes into near–genome-wide methylation maps, potentially enabling larger effective sample sizes at genome-wide resolution for downstream analyses.

ARUNA’s flexible framework presents several opportunities for further improvement. First, the current model and evaluations do not consider sequencing read depth, a key metric of uncertainty in beta measurements. In our experiments, no read-depth thresholding was applied to maintain maximal sequence resolution, but this may lead to training or evaluation on CpGs with high uncertainty. ARUNA’s future modifications may include ways to more directly incorporate this coverage information, either for better modeling or for joint uncertainty quantification in the upscaling. Second, long-range dependencies are captured only via stacked receptive fields of the deep convolutional architecture. Transformer models may prove powerful for this purpose by training with the slice setup, which allows for more data to stabilize training and provides opportunities for parallelization to improve scalability. Finally, interpretability remains an open opportunity: beyond performance metrics, future work could use targeted perturbations (masking, shuffling, or counterfactual edits of observed CpGs) to probe which local patterns and cross-sample signals drive predictions in specific genomic contexts (e.g., CGIs, promoters, or dynamic CpGs). We expect these directions to improve robustness and interpretability while maintaining ARUNA’s core advantage: a practical, scalable and accurate self-supervised approach to genome-wide imputation from sequencing assays without requiring paired measurements or extensive feature engineering.

