## Supplementary material for "ARUNA: Slice-based self-supervised imputation for upscaling DNA methylation sequencing assays"

### ARUNA: Supplementary materials

#### 1 Supplementary Tables

| Stage | Operation | Kernel | Stride | Channels (in $\rightarrow$ out) | Output shape |
|---|---|---|---|---|---|
| Input | Concat w/ positional embeds | – | – | $1 \rightarrow 2$ | $(B, 2, 8, 128)$ |
| Encoder | Conv2D + ReLU | $5 \times 5$ | $1 \times 1$ | $2 \rightarrow 32$ | $(B, 32, 8, 128)$ |
| | Conv2D + ReLU | $5 \times 5$ | $1 \times 1$ | $32 \rightarrow 64$ | $(B, 64, 8, 128)$ |
| | Conv2D + ReLU | $5 \times 5$ | $2 \times 2$ | $64 \rightarrow 128$ | $(B, 128, 4, 64)$ |
| | Conv2D + ReLU | $5 \times 5$ | $2 \times 2$ | $128 \rightarrow 128$ | $(B, 128, 2, 32)$ |
| | Conv2D | $5 \times 5$ | $2 \times 2$ | $128 \rightarrow 256$ | $(B, 256, 1, 16)$ |
| Bottleneck | Latent embedding | – | – | $256$ | $(B, 256, 1, 16)$ |
| Decoder | ConvTranspose2D + ReLU | $5 \times 5$ | $2 \times 2$ | $256 \rightarrow 128$ | $(B, 128, 2, 32)$ |
| | ConvTranspose2D + ReLU | $5 \times 5$ | $2 \times 2$ | $128 \rightarrow 128$ | $(B, 128, 4, 64)$ |
| | ConvTranspose2D + ReLU | $5 \times 5$ | $2 \times 2$ | $128 \rightarrow 64$ | $(B, 64, 8, 128)$ |
| | ConvTranspose2D + ReLU | $5 \times 5$ | $1 \times 1$ | $64 \rightarrow 32$ | $(B, 32, 8, 128)$ |
| Output | ConvTranspose2D + Sigmoid | $5 \times 5$ | $1 \times 1$ | $32 \rightarrow 1$ | $(B, 1, 8, 128)$ |

**Supplementary Table 1:** Detailed architecture for the symmetric pyramidal denoising convolutional autoencoder used in all experiments. Total parameters: 2.9M. Models were trained using the Adam optimizer with a learning rate of $5 \times 10^{-4}$ and early stopping. 10 repeats per target patch were used during training and 5 during inference.

#### 2 Supplementary Figures

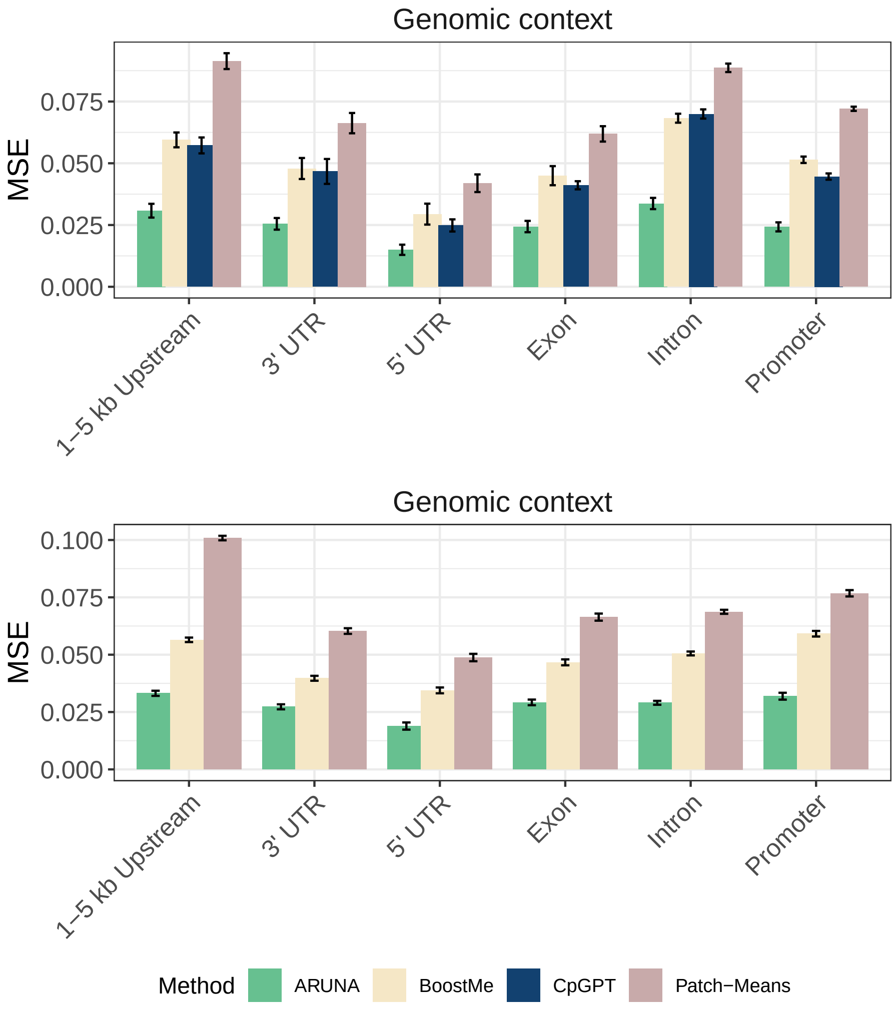


**Supplementary Figure 1:** Performance evaluations for the donor-holdout simulation setup in several genic contexts. Training and evaluation was performed using GTEx data with simulated missingness (mcar at 90% for training ARUNA and rrbs-like simulation for evaluations). Figures show MSE over held-out donor samples across aggregated over multiple folds and seeds. (Top) FullIntersect (2,985 ± 423 CpGs); (Bottom) GenomeScale (366,887 ± 4575 CpGs).

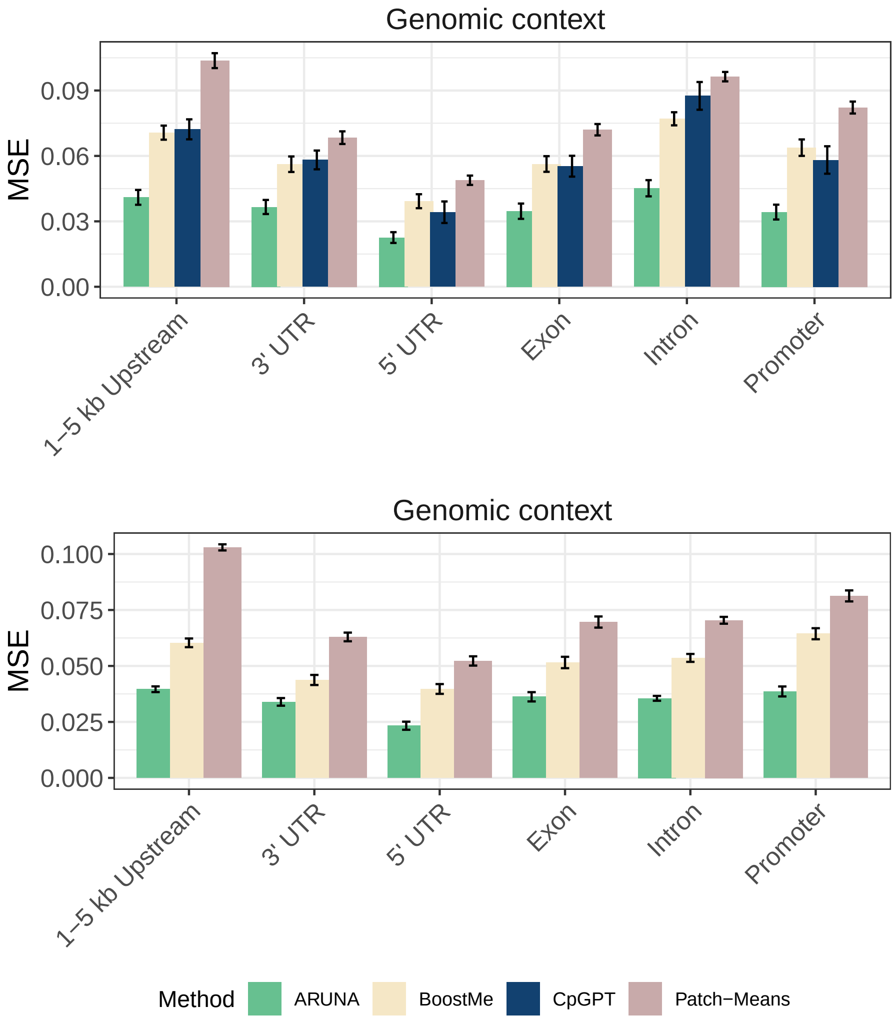


**Supplementary Figure 2:** Performance evaluations for the tissue-holdout simulation setup in several genic contexts. Generalization of models to unseen tissue types was assessed with MSE computed over the held-out tissue folds and seeds. (Top) FullIntersect (4,242 ± 1051 CpGs); (Bottom) GenomeScale (364,638 ± 5108 CpGs).
